# Single-cell RNA-seq reveals alterations in peripheral *CX3CR1* and nonclassical monocytes in familial tauopathy

**DOI:** 10.1101/2022.10.28.514304

**Authors:** Daniel W. Sirkis, Caroline Warly Solsberg, Taylor P. Johnson, Luke W. Bonham, Virginia E. Sturm, Suzee E. Lee, Katherine P. Rankin, Howard J. Rosen, Adam L. Boxer, William W. Seeley, Bruce L. Miller, Ethan G. Geier, Jennifer S. Yokoyama

## Abstract

**Background:** Emerging evidence from mouse models is beginning to elucidate the brain’s immune response to tau pathology, but little is known about the nature of this response in humans. In addition, it remains unclear to what extent tau pathology and the local inflammatory response within the brain influence the broader immune system.

**Methods:** To address these questions, we performed single-cell RNA sequencing (scRNA-seq) of peripheral blood mononuclear cells (PBMCs) from carriers of pathogenic variants in *MAPT*, the gene encoding tau.

**Results:** Analysis of ∼181,000 individual PBMC transcriptomes from *MAPT* pathogenic variant carriers (*n* = 8) and healthy non-carrier controls (*n* = 8) demonstrated striking differential expression in monocytes and natural killer (NK) cells. We observed a marked reduction in the expression of *CX3CR1* – the gene encoding the fractalkine receptor that is known to modulate tau pathology in mouse models – in monocytes and NK cells. We also observed a significant reduction in the abundance of nonclassical monocytes and dysregulated expression of nonclassical monocyte marker genes, including *FCGR3A*. Finally, we identified reductions in *TMEM176A* and *TMEM176B*, genes thought to be involved in the inflammatory response in human microglia. We confirmed differential expression of select biologically relevant genes dysregulated in our scRNA-seq data using droplet digital PCR as an orthogonal technique for quantitative validation.

**Conclusions:** Our results suggest that human peripheral immune cell expression and abundance are modulated by tau-associated pathophysiologic changes. *CX3CR1* and nonclassical monocytes in particular will be a focus of future work exploring the role of these peripheral signals in additional tau-associated neurodegenerative diseases.

## Background

Nearly 25 years after the discovery of pathogenic variants in *MAPT* (encoding the microtubule-associated protein tau) in familial frontotemporal dementia (FTD; [1,2]; reviewed in [3]), there are still no effective therapeutics capable of halting or delaying tau-associated neurodegeneration [4,5]. Diverse tau proteinopathies (tauopathies) also occur sporadically (i.e., in the absence of *MAPT* or other pathogenic variants) and are subdivided into primary tauopathies – which collectively fall under the umbrella term, frontotemporal lobar degeneration (FTLD)-tau – and secondary tauopathies, the most prominent example of which is Alzheimer’s disease (AD). Although much effort has gone into characterizing the natural history and longitudinal declines of *MAPT* pathogenic variant carriers (reviewed in [6]), we understand relatively little about the molecular mechanisms that impart risk for sporadic forms of tauopathy – whether primary or secondary.

The last decade has witnessed a major revival in interest in immune mechanisms that may modulate risk for neurodegeneration, with a primary focus on microglia, the parenchymal macrophages of the brain (reviewed in [7]). Significantly less progress has been made in elucidating peripheral blood or cerebrospinal fluid (CSF) leukocyte perturbations in – and responses to – neurodegeneration, although this is beginning to change (reviewed in [8–10]). For example, we now know of changes to CD8^+^ T cells in AD and CD4^+^ T cells in Lewy body dementia [11,12]. In addition, altered phospholipase C-ɣ2 signaling in peripheral lymphocytes has been reported in AD [13]. Beyond lymphocytes, changes in peripheral monocytes (particularly nonclassical [NC] monocytes) have been observed in Parkinson’s disease (PD) [14] and amyotrophic lateral sclerosis (ALS; [15,16]). In addition, patients with the hereditary white-matter disorder, adult-onset leukoencephalopathy with axonal spheroids and pigmented glia (ALSP), show striking reductions in peripheral NC monocytes [17]. Given that ALSP is thought to be driven by primary microglial defects (reviewed in [18]), the reduction in peripheral NC monocytes in this disorder suggests shared biology between these two cell types.

High-quality fluid biomarkers are being developed for AD (reviewed in [19,20]) and FTD (reviewed in [21]), but those that can distinguish between the major neuropathologic division within FTD (i.e., FTLD-tau vs. FTLD due to TDP-43 pathology [FTLD-TDP]) have been lacking [21] until very recently [22]. We reasoned that an in-depth exploration of peripheral immune dysregulation in tauopathy may reveal novel, blood-based biomarkers associated with FTLD-tau. Therefore, in an effort to define the peripheral immune signatures of tauopathy, we carried out single-cell RNA sequencing (scRNA-seq) of peripheral blood mononuclear cells (PBMCs) in individuals with pathogenic *MAPT* variants – who have (or will develop) FTLD-tau pathology – and cognitively normal, non-carrier controls. We identified striking changes in NC monocytes – both in terms of cellular abundance and gene expression – as well as natural killer (NK) cells and other cell types. Moreover, we identified *CX3CR1* as a potentially novel peripheral marker of tauopathy, suggesting parallel changes in *CX3CR1* in microglia and peripheral leukocytes in the course of tau-mediated neurodegeneration. We also identified additional candidate genes whose expression may be altered in the periphery in tauopathy (e.g., *FCGR3A* and *TMEM176A*/*B*). Considering recent findings in ALS and PD, and given that NC monocytes can be detected in the human brain [23], our results add to the weight of evidence suggesting the importance of NC monocytes across a spectrum of neurodegenerative diseases. Taken together, our results indicate that PBMCs represent an accessible and informative tissue source not only for biomarker discovery but also for elucidation of peripheral immune responses in the context of tauopathy.

## Methods

### Study Participants

All participants or their surrogates provided written informed consent prior to study participation; all aspects of the studies described here were approved by the University of California, San Francisco (UCSF) institutional review board. Sixteen individuals (*n* = 8 *MAPT* pathogenic variant carriers and *n* = 8 cognitively normal, non-carrier controls) participated in this study. Individuals were recruited as part of ongoing studies of FTD and related neurodegenerative diseases at the UCSF Memory and Aging Center (MAC). *MAPT* pathogenic variant carriers had clinical syndromes of bvFTD (*n* = 4), frontal AD (*n* = 1), had subjective cognitive impairment (*n* = 1), or were considered clinically normal (*n* = 2); *MAPT* variant carriers diagnosed as clinically normal were considered to be presymptomatic (*n* = 2). Pathogenic *MAPT* variants represented within this study were p.P301L (*n* = 1), p.S305I (*n* = 1), p.S305S (*n* = 1), p.R406W (*n* = 3), and IVS10+16 (*n* = 2). Pathogenic variant carriers and cognitively normal controls were sex-matched, and equal numbers of female and male participants were included in the cohort. Age was not significantly different between carrier and control groups, as assessed using unpaired t-test. Demographic information for study participants is included in Table 1.

**Table 1.** Demographic characteristics of cohort.

|  | Control | *MAPT* variant carrier |
| --- | --- | --- |
| *n* | 8 | 8 |
| Age, mean (SD) | 52.8 (10.0) | 54.4 (11.4) |
| Sex, *n* female | 4 | 4 |
| Clinical syndrome (*n*) | clinically normal (8) | bvFTD (4), frontal AD (1), subjective cognitive impairment (1), presymp. (2) |
| *MAPT* variants (*n*) | N/A | p.P301L (1), p.S305I (1), p.S305S (1), p.R406W (3), IVS10+16 (2) |
| Splicing variants, *n* | N/A | 4 |
| CDR-SB, mean (SD) | 0.0 (0.0) | 5.8 (5.7)<br>7.8 (5.3)<sup>#</sup><br><sup>#</sup>symptomatic carriers only |

Abbreviations: AD, Alzheimer's disease; bvFTD, behavioral variant frontotemporal dementia; CDR-SB, Clinical Dementia Rating scale Sum of Boxes; presymp., presymptomatic; SD, standard deviation.

### Clinical Assessment

Study participants underwent a multistep screening process prior to an in-person clinical evaluation at the UCSF MAC, which included a neurological exam, cognitive assessment, and medical history [24]. Each participant’s study partner was interviewed to assess the participant’s functional abilities. A multidisciplinary team composed of a behavioral neurologist, a neuropsychologist, and a registered nurse established clinical diagnoses for cases according to consensus criteria for FTD and its subtypes [25] or frontal AD [26]. Controls and presymptomatic *MAPT* carriers had a normal neurological exam and a Clinical Dementia Rating scale Sum of Boxes (CDR-SB) [27] score of 0. All controls screened negative for disease-causing pathogenic variation in established AD and FTD genes.

### Cell Isolation

Human PBMCs were obtained from study participants at the UCSF MAC. Blood specimens were collected in yellow-top acid-citrate-dextrose vacutainer tubes (BD Biosciences) and processed within five hours of collection. PBMCs were isolated via Ficoll density gradient centrifugation using Lymphosep separation medium (MP Biomedicals), washed with Ca^2+^- and Mg^2+^-free PBS (ThermoFisher), and treated with one application of red blood cell lysis buffer (Biolegend). After a final wash step in PBS, PBMCs were diluted to a density of 1.5 × 10^6^ cells/ml in freezing media composed of 10% DMSO in FBS and immediately frozen at -80°C. After two weeks, samples were transferred to liquid nitrogen for long-term storage.

### Single-Cell RNA-seq

PBMCs were thawed and prepared for scRNA-seq using the Chromium Single Cell 3’ v2 kit according to the manufacturer’s instructions (10x Genomics). Samples were processed in two separate batches of eight samples each, with four *MAPT* variant carriers and four controls included in each batch. To further minimize the potential for batch effects, each batch contained equal numbers of samples from female and male participants. After sample thawing, counting, and dilution, PBMCs underwent standard 10x processing, 3’ gene expression library construction steps, and next-generation sequencing at the UCSF Genomics CoLab and Institute for Human Genetics (IHG).

### Sequencing Data Processing

For each of the two batches, single-cell 3’ libraries generated from eight samples were pooled and sequenced on one lane of a NovaSeq S4 flow cell. Raw sequencing reads were aligned to GRCh38, and feature-barcode matrices were generated using Cell Ranger version 3.0.2.

### Quality Control and Clustering

We obtained a total of 7.3 × 10^9^ reads and detected ∼214,000 cells across the two independent 10x and sequencing batches, yielding a moderate sequencing depth [28,29] of ∼34,000 mean reads/cell. We detected ∼3,700 median UMI counts/cell and ∼1,100 median genes/cell (Table S1). There were no significant differences in the number of cells captured per sample, the number of reads per sample, or the mean read depth per sample when comparing the *MAPT* pathogenic variant carrier group to the non-carrier control group. Subsequent quality-control (QC) and downstream analysis steps were performed using Seurat v4.1 [30,31]. QC filtering was applied to individual-sample feature-barcode matrices and consisted of the following steps: (i) genes detected in < 10 cells were removed; (ii) cells with ≤ 500 detected genes were removed; (iii) cells with ≤ 500 counts and those with ≥ 20,000 counts were removed; (iv) cells with mitochondrial mapping percentages ≥ 10 were removed; (v) doublets were identified and removed using DoubletFinder v2.0.3 [32,33] using the recommended parameter settings. After stringent QC filtering, ∼181,000 cells remained for downstream analysis (Table S2).

After QC, we performed the following additional processing steps: (i) we applied sctransform [34]–a method for scRNA-seq count normalization and variance stabilization–at the individual-sample level, including mitochondrial mapping percentage as a covariate [34,35], to minimize variability due to differences in sequencing depth between samples; (ii) the 16 individual samples were integrated with `FindIntegrationAnchors` and `IntegrateData`, specifying ‘sctransform’ as the normalization method and reciprocal principal component analysis (PCA) as the reduction. Subsequently, PCA was performed followed by uniform manifold approximation and projection (UMAP) reduction using the first 30 PCs; clustering was performed using a resolution parameter of 0.5. This resulted in the generation of 21 distinct clusters that were annotated on the basis of marker gene expression, identified using `FindMarkers` and literature searches.

### Differential Expression Analysis

Differential expression analysis was performed using limma [36–39] on individual clusters and sctransform-normalized data, with *MAPT* pathogenic variant carrier status as the contrast and age, sex, and scRNA-seq batch as covariates. To account for multiple testing, a false discovery rate-corrected *p*-value (*p*_FDR_) < 0.05 was considered statistically significant. For visualization of selected differentially expressed genes (DEGs), we used violin plots displaying normalized count data generated via Seurat’s `NormalizeData` function.

### Cluster Proportionality

Cluster proportions were determined for individual samples by dividing the number of cells in a given cluster by the total number of cells in all clusters (after QC filtering) for each individual. Differences in cluster proportionality were assessed visually for all clusters according to *MAPT* variant carrier status. Only the NC monocyte cluster (cluster 11) showed a clear difference in proportionality between carriers and controls. Significance for cluster 11 proportionality, as well as NC monocyte subcluster ratios, was determined by linear model, covarying for age, sex, and scRNA-seq batch.

### STRING Network Analysis

DEGs with *p*_FDR_ < 0.05 and log_2_ fold-changes (LFC) > 0.1 for selected cell clusters were submitted for analysis via the STRING database (v11.5) [40] using the following parameters: the full STRING network was queried; network edge thickness indicated the confidence of the interaction; active interaction sources included databases, experiments, and text mining; a minimum interaction score of 0.4 was required; only query genes/proteins were displayed; disconnected nodes (i.e., DEGs with no interaction partners) were not displayed; gene modules were color-coded according to the results of Markov cluster algorithm (MCL) clustering [41]. For clarity, within the Results section we refer to specific MCL clusters (containing DEGs) as gene/protein ‘modules,’ while reserving the term ‘cluster’ to refer to cell clusters generated via scRNA-seq analysis.

### Analysis of Mouse RNA-seq Data

Publicly available mouse RNA-seq data were downloaded from GEO (accession GSE93180) and originally published in [42]. Briefly, hippocampal microglia (Cd11b^+^ myeloid cells) were sorted from 6-month-old male hMAPT-P301S transgenic mice (*n* = 6) or non-transgenic littermates (*n* = 6), then RNA was extracted and processed for RNA-seq. We reanalyzed the available count data using DESeq2 [43] and plotted normalized counts for mouse *Cx3cr1* using ggplot2 [44].

### RNA Extraction

To minimize biological variability and facilitate orthogonal validation, for droplet digital (dd)PCR experiments, we used PBMCs isolated from 15 of the 16 participants originally selected for scRNA-seq analysis. One additional cognitively normal control sample was used for ddPCR due to unavailability of one control sample used for scRNA-seq. RNA was extracted from the PBMCs using the RNeasy Micro Kit (Qiagen) and isolated RNA was quantified and its quality was assessed using the RNA 6000 Pico Bioanalyzer kit (Agilent). PBMC RNA samples had RNA integrity number (RIN) values ranging from 9.2–9.9, indicating high-quality RNA [45].

### Droplet Digital PCR

One ng of total RNA was used for single-tube reverse transcription (RT) and ddPCR using the One-Step RT-ddPCR Advanced kit (Bio-Rad). Droplets were generated and subsequently analyzed using the QX100 system (Bio-Rad) at the UCSF Center for Advanced Technology (CAT). Reactions were prepared and run essentially according to the manufacturer’s instructions. For steps in which a temperature range was specified, we used the following parameters: RT was performed at 50°C, annealing/extension occurred at 55°C, and samples were held at 12°C in the C1000 thermocycler (Bio-Rad) prior to analysis on the droplet reader. To confirm specificity, we ran the following control reactions: wells lacking RNA but containing all other components and wells lacking reverse transcriptase but containing all other components. PrimePCR ddPCR Gene Expression primer–probe mixes coupled to FAM or HEX (Bio-Rad) were used to amplify specific genes.

### Additional Statistical Analysis

For the analysis of ddPCR data, we performed linear modeling in R to assess whether *MAPT* carrier status predicted differences in gene expression while covarying for age and sex. Log_2_-transformed absolute concentration data for *CX3CR1, FCGR3A, TMEM176A/B*, and *C3AR1* (or the ratios of these values with those of reference gene *EEF2*) were used for analyses assuming data normality, while non-transformed data are displayed in the plots. Plots were generated with ggplot2.

## Results

After QC filtering, clustering of ∼181,000 PBMCs generated 21 primary clusters representing the major lymphoid and myeloid cell types, including CD4^+^ and CD8^+^ T cells, B cells, NK cells, monocytes, and dendritic cells (Fig. 1A; Fig. S1). As expected, and in contrast to many other FTD-associated genes, *MAPT* expression was barely detectable in PBMCs (Fig. S2). We reasoned, therefore, that any changes detected in PBMC cell-type proportionality or gene expression in *MAPT* pathogenic variant carriers would most likely represent a response to tau neuropathology or neurodegeneration rather than cell-intrinsic changes occurring directly downstream of variant *MAPT*. Of all PBMC clusters, only one showed a clear, consistent change in abundance in *MAPT* pathogenic variant carriers relative to controls. This cluster (11), which represents *FCGR3A*^+^ (CD16^+^) NC monocytes (Fig. 1A, B), localized in UMAP space near the more abundant *CD14*^+^ classical monocyte cluster (2) and the *CLEC10A*^+^ conventional dendritic cell (cDC) cluster (14). In particular, NC monocytes, expressed as a percentage of total PBMCs for each participant, were significantly reduced in *MAPT* carriers (Fig. 1C). To gain more fine-grained insight into the nature of this change in abundance, we subsetted and re-clustered myeloid clusters 2, 11, and 14. Assessing the myeloid subclusters (Fig. 1D, left), we discovered that normalizing the NC monocyte subcluster to the *CLEC10A*^+^, *CD1C*^+^ cDC subcluster (corresponding to the most abundant blood cDC population, known as cDC2 [46]; Fig. 1D, lower right panel) yielded better separation between *MAPT* variant carriers and non-carrier controls (Fig. 1E). Further assessing the myeloid subclusters, we found that normalizing NC monocyte numbers to total cDCs (i.e., cDC1 + cDC2) gave a similar result (Fig. S3A). On the other hand, expressing NC monocyte abundance as a fraction of all monocytes or all myeloid cells did not significantly differentiate carriers from controls (Fig. S3B, C).

**Figure 1.**
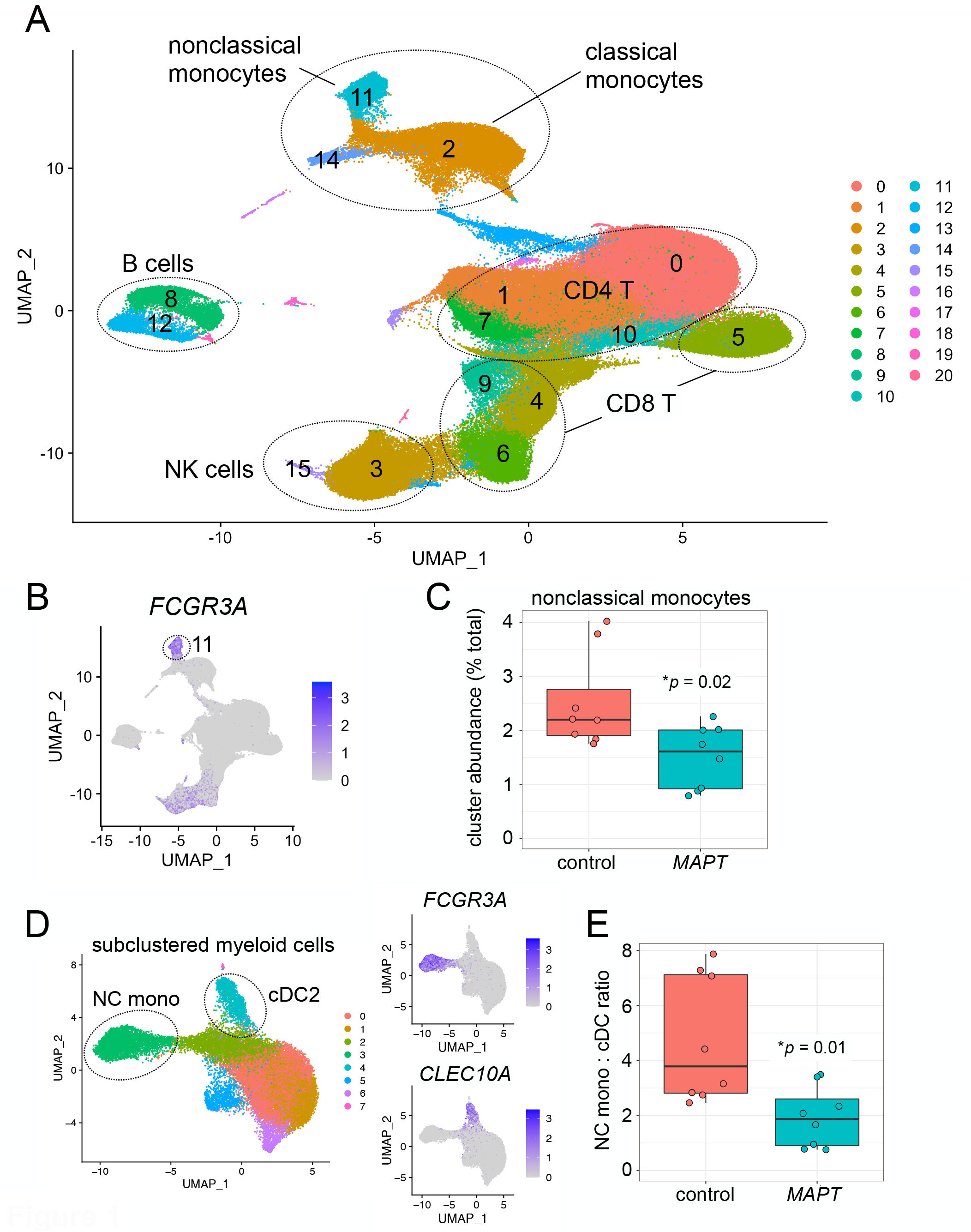
Single-cell RNA-seq reveals reductions in nonclassical monocytes in *MAPT* pathogenic variant carriers. **A** Two-dimensional UMAP plot of ∼181,000 PBMCs from *MAPT* variant carriers and non-carrier controls, colored by cluster identity. Major cell types are labeled within the plot. **B, C** Cluster 11, marked by high *FCGR3A* expression and identified as NC monocytes, was significantly reduced in *MAPT* carriers (*p* = 0.02; data are expressed as percentage of total PBMCs for each sample). **D** Myeloid cells (clusters 2, 11, 14) were subset and re-clustered. NC monocyte and cDC2 subclusters were identified by *FCGR3A* and *CLEC10A* expression, respectively (**D**, right). **E** The ratio of NC monocytes to cDC2 was significantly reduced in *MAPT* variant carriers (*p* = 0.01).

We next performed differential expression analysis on each of the primary cell clusters, comparing *MAPT* variant carriers to non-carrier controls and adjusting for age, sex, and scRNA-seq batch. Focusing initially on DEGs with absolute LFC > 0.2, we determined that cDC, NC monocyte, and NK cell clusters had the highest number of DEGs (Fig. 2A, B). We further characterized the DEGs of NC monocytes and NK cells by querying the STRING database [40] for functional and physical interactions, now using a more-permissive LFC cutoff of 0.1 to facilitate population of the gene interaction network (see Table S3 for DEG lists for all clusters).

**Figure 2.**
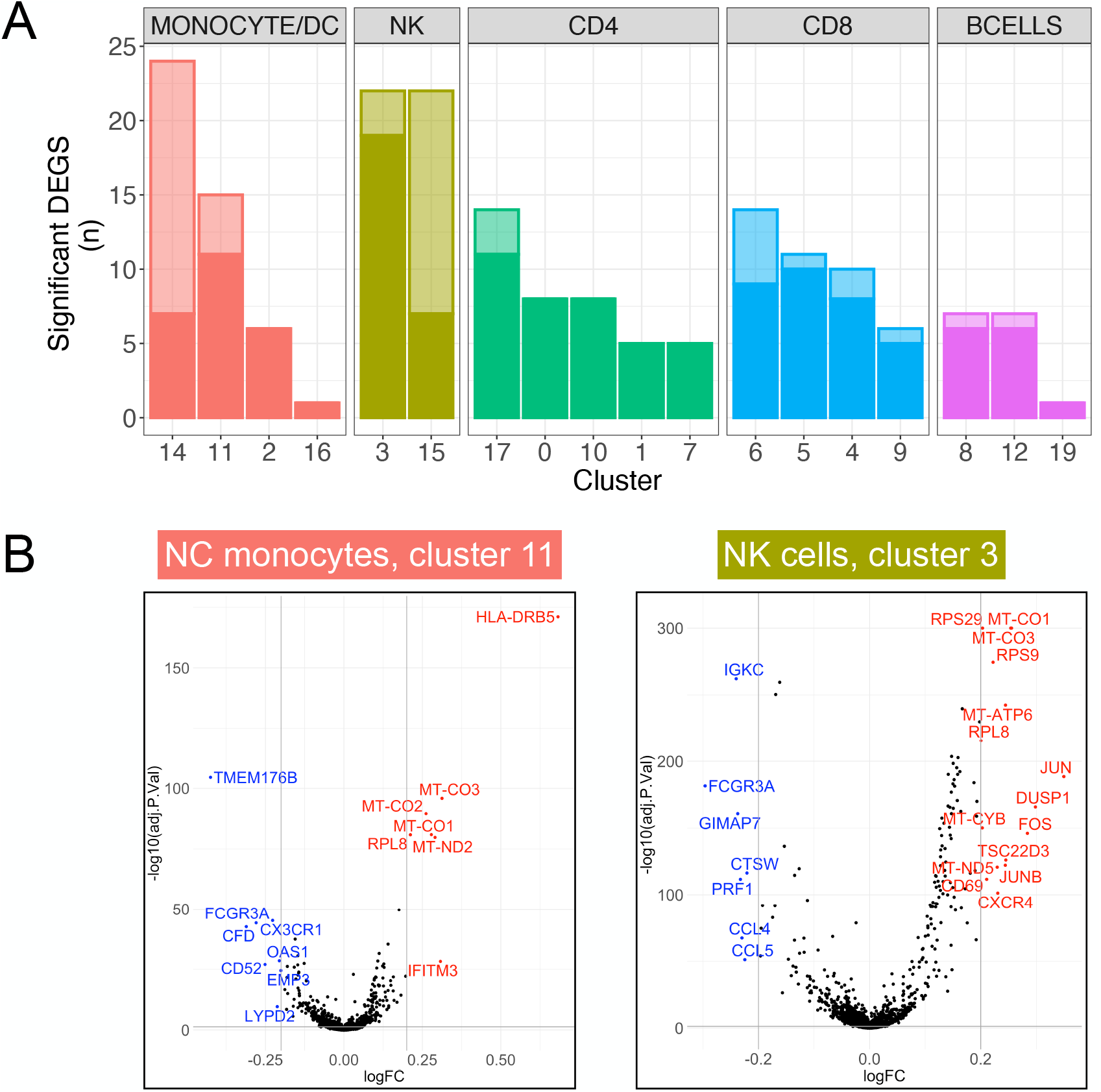
Differential expression in *MAPT* variant carriers by cell cluster. **A** Clusters are grouped by cell type and ranked by the number of DEGs with *p*_*FDR*_ < 0.05 and absolute LFC > 0.2. Differential expression was determined in *MAPT* variant carriers relative to non-carrier controls while covarying for age, sex, and scRNA-seq batch. Solid-colored portions of the bars indicate DEGs shared by at least one other cluster, while the translucent portions indicate DEGs unique to a given cluster. cDCs (cluster 14), NK cells (clusters 3 and 15), and NC monocytes (cluster 11) had the highest numbers of DEGs with absolute LFC > 0.2. **B** Volcano plots of the NC monocyte cluster and major NK cell cluster; DEGs with absolute LFC > 0.2 are labeled in blue (down-regulated) or red (up-regulated). Several NK cell DEGs (right) with -log_10_(*p*_FDR_) values > 300 were set to 300 for visualization purposes.

*MAPT* carriers showed striking up-regulation of many ribosomal and mitochondrial genes that, respectively, formed large interaction modules (Fig. 3A, B). The biological significance of these coordinately up-regulated DEGs is unclear, but tau is known to interact with and affect multiple ribosomal subunits [47–50] and mitochondrial proteins [50,51]. Because the identified mitochondrial DEGs represent a subset of the mitochondrial genes used during QC (see Methods) to filter out putatively damaged cells [35], we considered the possibility that – despite the removal of cells with high mitochondrial mapping percentages (≥ 10%) – the apparent up-regulation of mitochondrial DEGs may nevertheless be associated with higher mitochondrial mapping percentage in *MAPT* variant carriers. Consistent with this possibility, mitochondrial mapping percentage was subtly but significantly higher in *MAPT* carriers in the NC monocyte and NK cell clusters (11 and 3, respectively; Fig. S4A, B). On the other hand, mitochondrial DEGs were absent from the cDC cluster (14; Fig. S4D; Table S3) – despite this cluster having the highest overall number of DEGs with LFC > 0.2 (Fig. 2A) – and mitochondrial mapping percentage was not significantly increased in variant carriers in this cluster (Fig. S4C). This suggests (i) cell-type-specific and, presumably, biologically relevant dysregulation of mitochondrial genes in *MAPT* variant carriers, consistent with [50]; and (ii) that the up-regulation of mitochondrial DEGs in the NC monocyte and NK cell clusters is associated with increased mitochondrial mapping percentage. To test the latter possibility, we next included mitochondrial mapping percentage as an additional covariate in the differential expression analyses, and, as expected, nearly all mitochondrial DEGs that previously had estimated LFCs > 0.1 no longer passed this threshold (not shown). Thus, the presence of mitochondrial DEGs is associated with increased mitochondrial mapping percentage. Importantly, coordinated up-regulation of mitochondrial genes could lead to subtle shifts in mitochondrial mapping percentage, and therefore the causality underlying this relationship is unclear.

**Figure 3.**
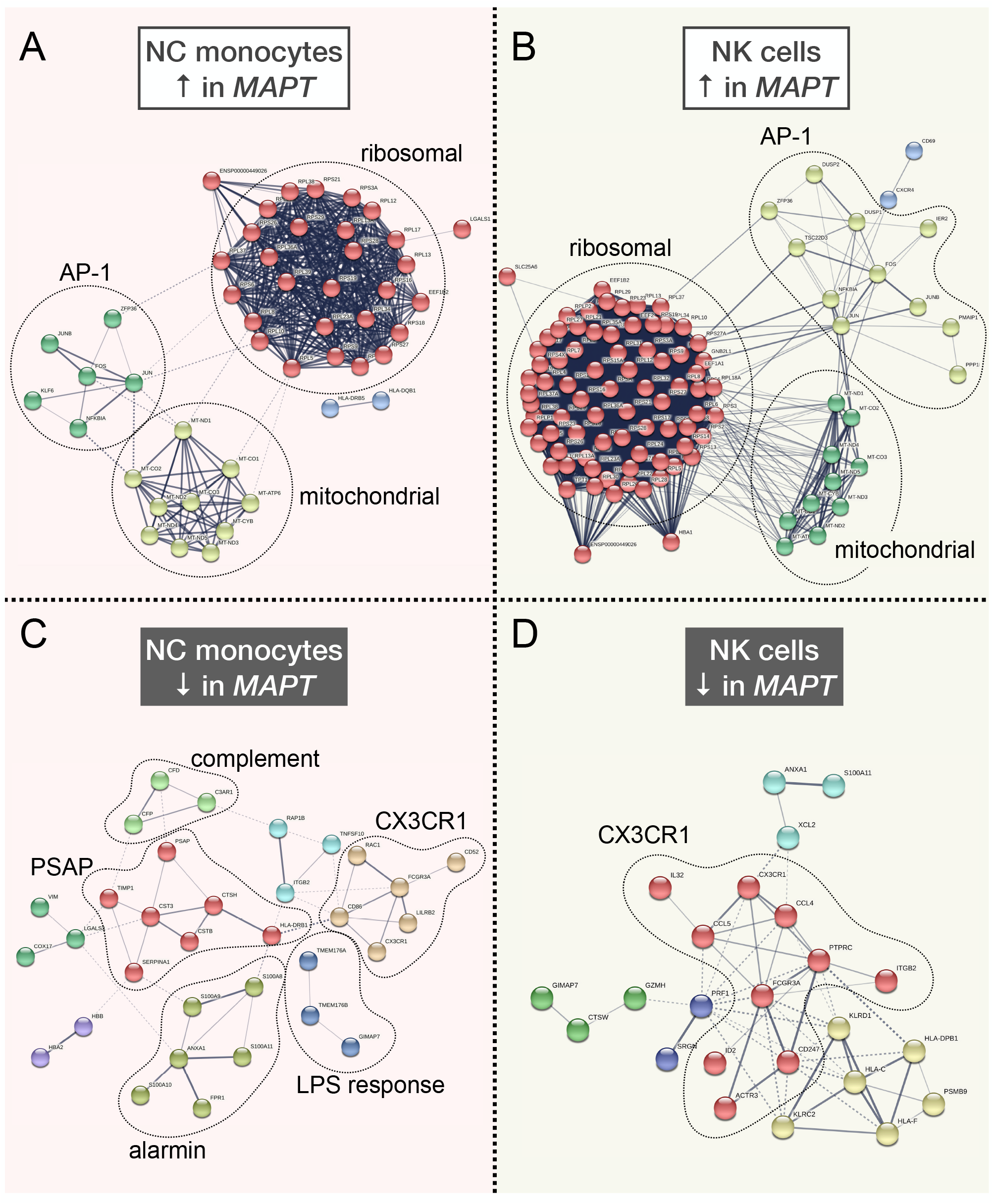
STRING interaction networks reveal relationships among nonclassical monocyte and natural killer cell differentially expressed genes. **A, B** Up-regulated DEGs in NC monocytes and NK cells had similar overall network architecture, with large ribosomal and mitochondrial modules, and a third module containing members of the AP-1 transcription factor, among other genes. **C** The down-regulated DEGs in NC monocytes contained a module featuring *CX3CR1* and *FCGR3A* as members, in addition to modules harboring genes involved in LPS response, the alternative and classical complement cascades, the *S100* alarmin molecules, as well as *PSAP*. **D** Down-regulated DEGs in NK cells also featured a large module featuring both *CX3CR1* and *FCGR3A*. All DEGs with *p*_*FDR*_ < 0.05 and absolute LFC > 0.1 from clusters 3 and 11 were input into the STRING database as described in the Methods section. Modules are colored according to the results of MCL clustering.

The other major up-regulated STRING module found in both NC monocytes and NK cells contained members of the AP-1 transcription factor family [52], including *FOS, JUN*, and *JUNB* (Fig 3A, B). Multiple members of this module (*FOS, DUSP1*) were previously found to be up-regulated via bulk measurements of both PBMCs and hippocampus in AD [53], suggesting consistent dysregulation of AP-1 transcription factor genes in both primary and secondary tauopathies. *NFKBIA*, encoding the NF-κB inhibitor-a, also appears in this module in both NC monocytes and NK cells, and this gene is highly up-regulated by treatment of microglia with tau paired-helical filaments [54] and fibrils [55] and in the brain in late-onset AD [56].

In terms of significantly down-regulated genes, both NC monocytes and NK cells contained a module populated by *FCGR3A* and *CX3CR1* (Fig 3C, D). Although *FCGR3A* is an established marker of NC monocytes, it is also expressed by a subset of NK cells (Fig. 1B). *CX3CR1* is highly expressed by both NC monocytes and NK cells (Fig. 4A) and, strikingly, is a well known modulator of tau pathophysiology in the brain [57–59]. Additional down-regulated modules in NC monocytes included those containing components of the complement pathway (*CFD, CFP, C3AR1* [60,61]), members of the S100 alarmin family (*S100A8*-*11* [62,63]), highly lipopolysaccharide (LPS)-responsive microglial genes (*TMEM176A, TMEM176B* [64]), and the lysosomal gene *PSAP*, which promotes pro-inflammatory activity in monocytes [65] (Fig. 3C). Collectively, the down-regulation of these gene modules in *MAPT* pathogenic variant carriers suggests dampening of latent pro-inflammatory pathways in NC monocytes and therefore emergence of a heightened anti-inflammatory phenotype. Of note, this apparent phenotypic change also occurred within a diminished population of circulating NC monocytes.

**Figure 4.**
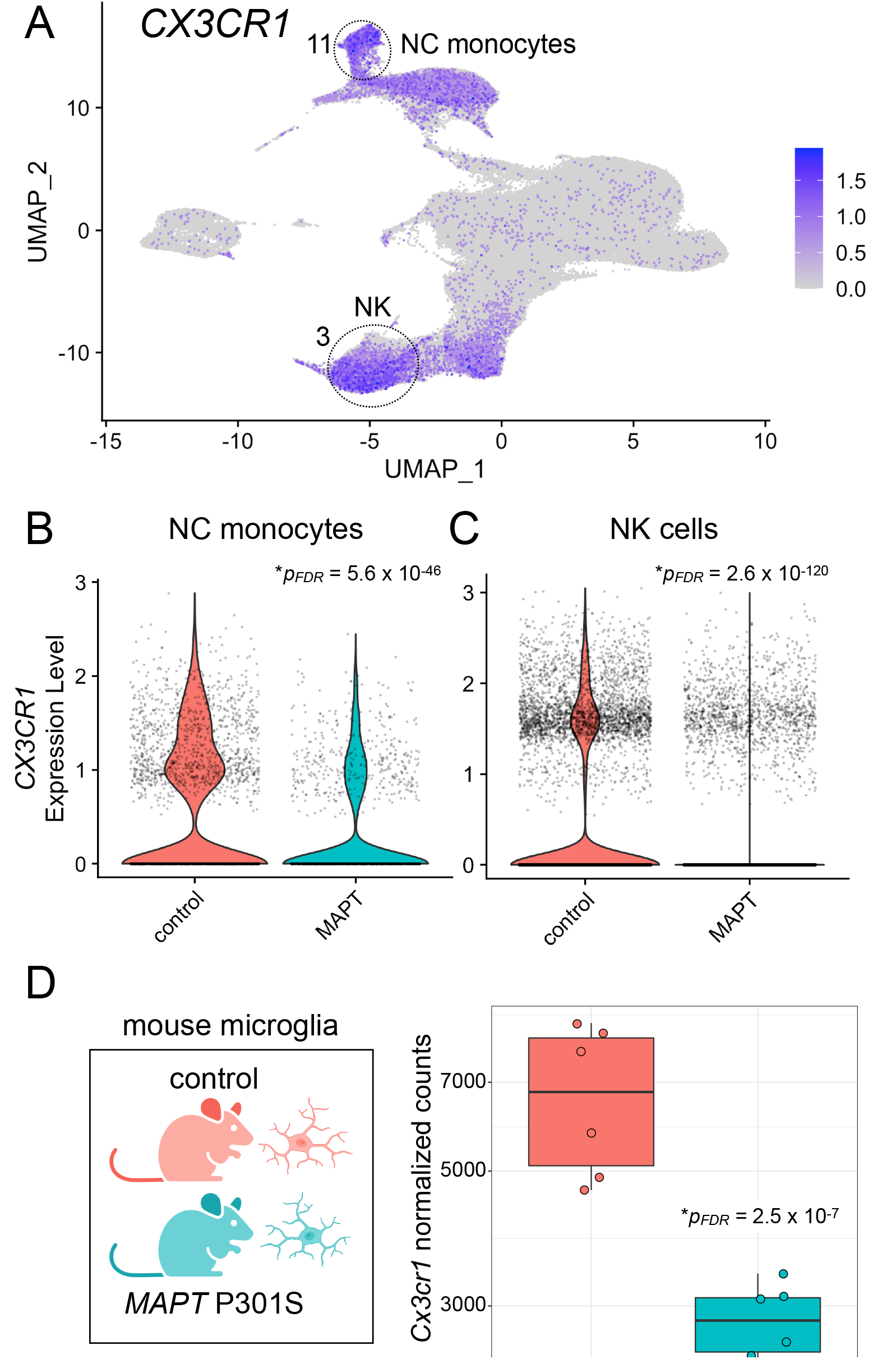
*CX3CR1* expression is reduced in peripheral myeloid and lymphoid cells in familial tauopathy. **A** *CX3CR1* is robustly expressed in both NC monocytes (cluster 11) and NK cells (cluster 3). NC monocytes (**B**; *p*_*FDR*_ = 5.6 × 10^−46^) and NK cells (**C**; *p*_*FDR*_ = 2.6 × 10^−120^) both show significantly reduced expression of *CX3CR1* in *MAPT* pathogenic variant carriers. **D** Reanalysis of publicly available bulk RNA-seq data from mouse hippocampal CD11b^+^ microglia demonstrated a significant reduction (*p*_*FDR*_ = 2.5 × 10^−7^) of *Cx3cr1* in the *MAPT* P301S model.

We next sought to validate a handful of DEGs which showed robust changes by differential expression analysis, often in multiple cell types, focusing on those deemed most likely to be biologically relevant to tau pathophysiology. In particular, we selected the following genes for validation via an orthogonal technique, ddPCR: *CX3CR1, FCGR3A, TMEM176A, TMEM176B*, and *C3AR1*. As noted above, *CX3CR1* has many well-established connections to tau pathology via its role in microglia. However, to our knowledge, its role in peripheral myeloid cells has not been studied in the context of neurodegenerative disease. *FCGR3A* not only serves as a marker gene for the significantly reduced population of NC monocytes but was also significantly down-regulated in both NC monocytes and NK cells of *MAPT* carriers. *TMEM176A*/*B* are less well known but have been shown to be extremely responsive to LPS treatment in human microglia [64], are homologs of the *MS4A* gene family involved in risk for AD [66], and have also been shown to be dysregulated in PBMCs from AD patients [53]. Finally, *C3AR1* has been implicated as a key player in the spread of tau neuropathology in mouse models [67], while the complement system more generally is thought to be a key regulator of neuronal loss in primary tauopathy as well as AD [68]. We first discuss validation of the *CX3CR1* finding below.

*CX3CR1* showed robust expression in NC monocytes and NK cells (Fig. 4A), and its expression was significantly reduced in both cell types in *MAPT* pathogenic variant carriers (Fig. 4B, C). To determine whether down-regulation in peripheral leukocytes was mirrored by changes in brain myeloid cells in the context of tauopathy, we analyzed a publicly available brain RNA-seq dataset [42] derived from *MAPT* P301S mouse hippocampal microglia. P301S microglia also displayed down-regulation of *Cx3cr1* (Fig. 4D), indicating (i) consistent *CX3CR1* dysregulation between microglia and peripheral immune cells in the context of tauopathy; and (ii) conserved *CX3CR1* expression changes between mouse and human in tauopathy. To validate the change in *CX3CR1* expression in peripheral immune cells, we isolated PBMC RNA from *MAPT* pathogenic variant carriers and non-carrier controls and performed ddPCR. ddPCR confirmed the reduction in *CX3CR1* (Fig. 5A). Normalizing the *CX3CR1* signal to reference gene *EEF2* gave similar results (Fig. S5A, B). When *MAPT* variant carriers were stratified by variant class (splicing vs. non-splicing), both groups showed significant reductions in *CX3CR1* (Fig. 5B), suggesting that multiple mechanistic forms of familial tauopathy converge on perturbation of *CX3CR1* expression.

**Figure 5.**
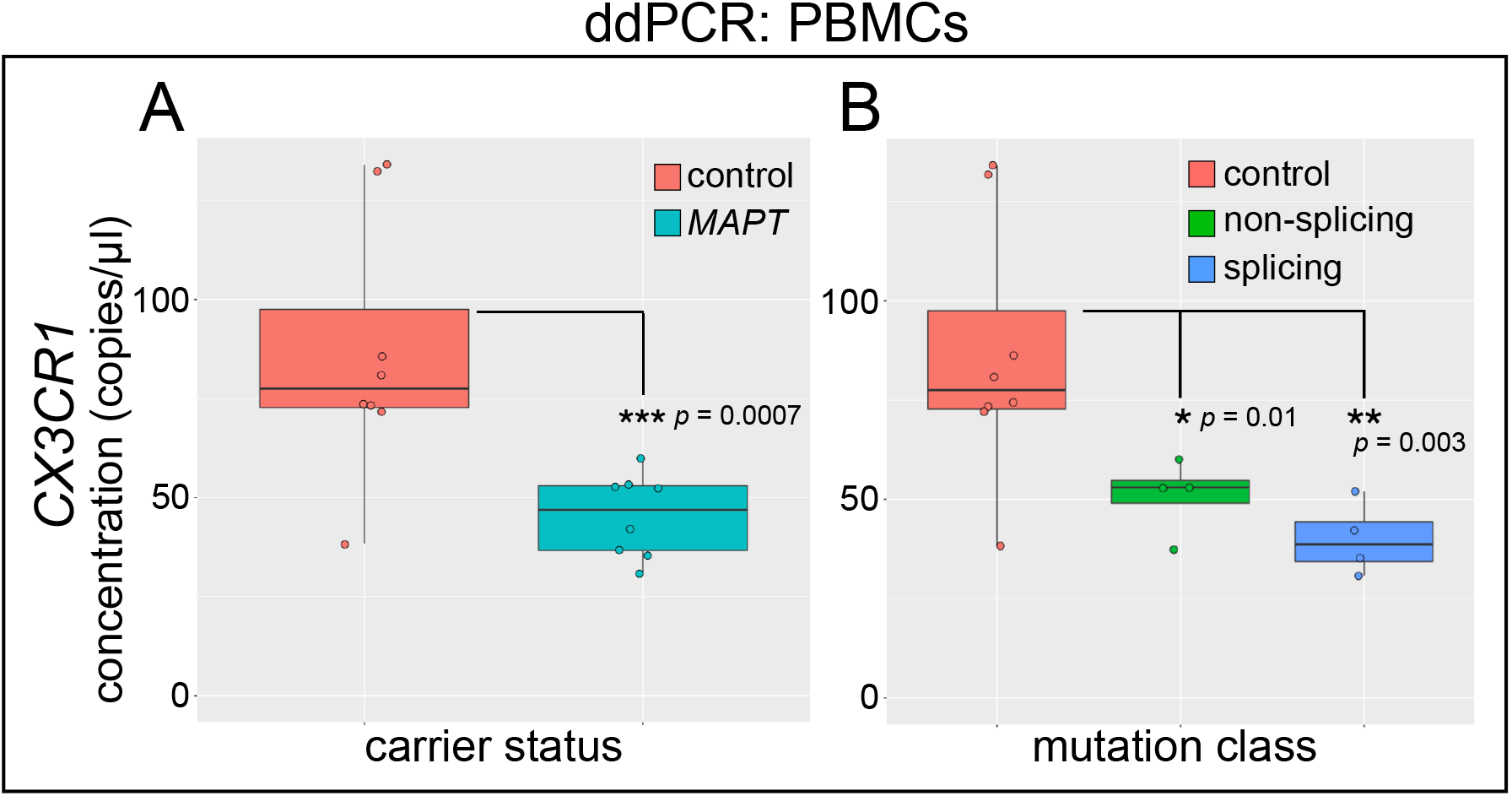
Confirmation of reduced *CX3CR1* expression in *MAPT* variant carrier PBMCs via ddPCR. RNA was isolated from PBMCs from *MAPT* variant carriers and healthy, non-carrier controls; gene expression was determined by RT-ddPCR. **A** *CX3CR1* was significantly reduced in *MAPT* carrier PBMCs (*p* = 0.0007) relative to controls. **B** Separation of samples according to *MAPT* variant class (non-splicing and splicing) reveals that *CX3CR1* was significantly reduced in both groups, relative to controls (non-splicing, *p* = 0.01; splicing, *p* = 0.003).

We next focused on validation of *FCGR3A*. We predicted that *FCGR3A* would show a robust decrease in expression in *MAPT* variant carriers in bulk PBMC RNA given that (i) NC monocytes are reduced in abundance in carriers (Fig. 1C), and (ii) *FCGR3A* expression was reduced in both NC monocytes and NK cells by scRNA-seq differential expression analyses (Fig. 6A, 2B). Indeed, ddPCR analysis showed a significant reduction in *FCGR3A* expression in *MAPT* pathogenic variant carrier PBMCs (Fig. 6B). The observed reduction likely reflects both reduced expression of *FCGR3A* and reduced levels of NC monocytes, which express *FCGR3A* at high levels. As with *CX3CR1*, normalization of the *FCGR3A* signal to reference gene *EEF2* did not affect the results (Fig. S5C). We also analyzed additional marker genes of the NC monocyte cluster, including *VMO1* and *IFITM3. VMO1*, which is specifically expressed in cluster 11 (consistent with [17]), and *IFITM3*, which is enriched in cluster 11, both showed significant differential expression in *MAPT* variant carriers (Fig. 6C, D; Table S3), indicating changes in multiple genes representing NC monocyte identity.

**Figure 6.**
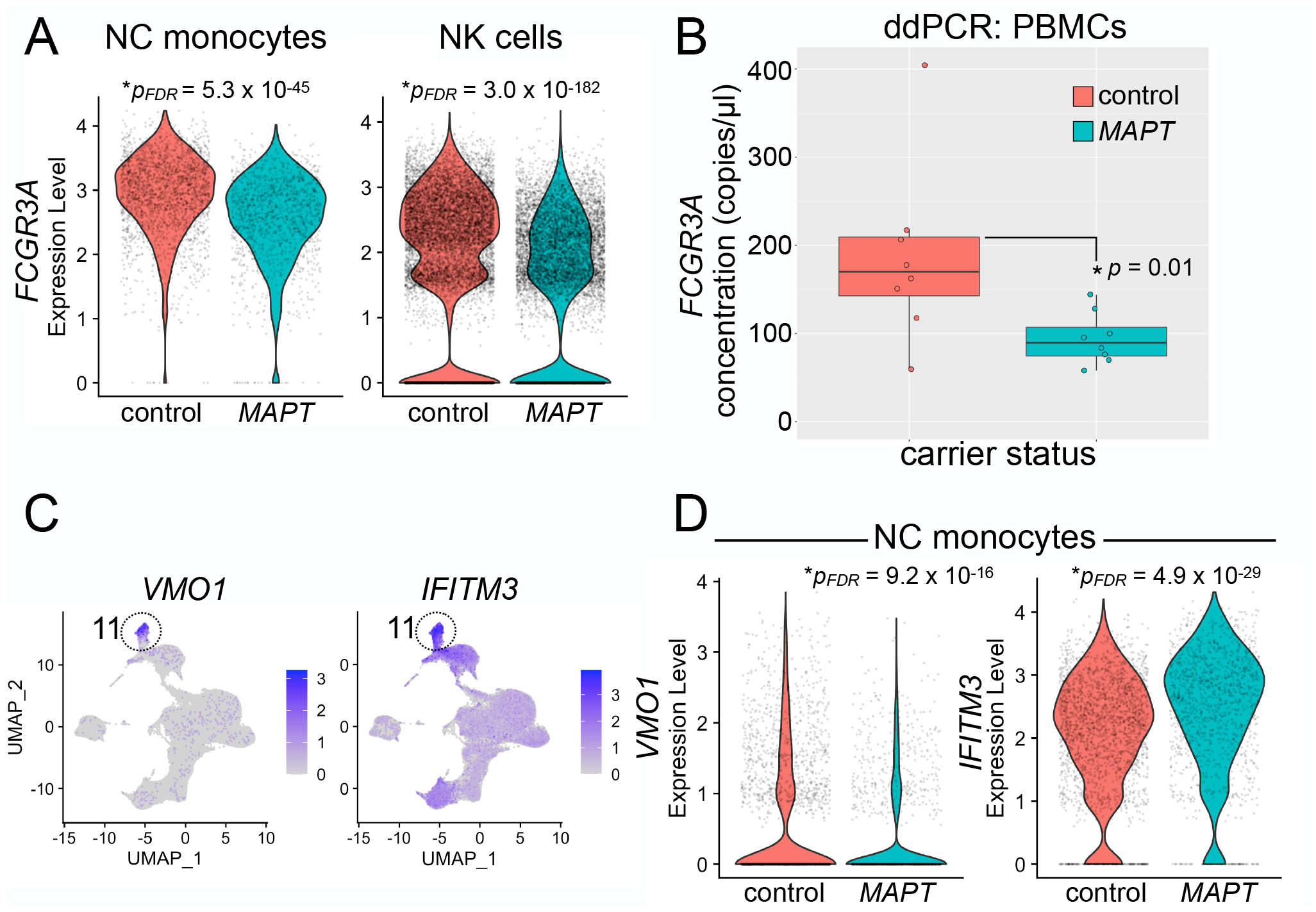
Analysis of nonclassical monocyte marker genes in *MAPT* variant carriers. **A** *FCGR3A*, the NC monocyte marker gene encoding CD16, is robustly expressed not only in NC monocytes but also in NK cells. *FCGR3A* is significantly reduced in both NC monocytes (left; *p*_*FDR*_ = 5.3 × 10^−45^) and NK cells (right; *p*_*FDR*_ = 3.0 × 10^−182^) in *MAPT* pathogenic variant carriers. **B** ddPCR confirmed a reduction in *FCGR3A* reduction in *MAPT* variant carrier PBMCs (p = 0.01). **C** Additional genes expressed specifically (*VMO1*, left) or enriched in (*IFITM3*, right) NC monocytes showed significant alterations (**D**) in *MAPT* variant carrier NC monocytes. **D** *VMO1* (left) was significantly reduced (*p*_*FDR*_ = 9.2 × 10^−16^), while *IFITM3* (right) was significantly increased (*p*_*FDR*_ = 4.9 × 10^−29^) in *MAPT* carriers.

*TMEM176A*/*B* represent poorly characterized genes of the extended *MS4A* family [66] thought to be involved in microglial LPS response [64], antigen presentation [69], and inflammasome regulation [70]. *TMEM176A*/*B* were highly expressed in classical and NC monocytes (clusters 2 and 11; Fig. 7A) and were strongly down-regulated in *MAPT* variant carrier NC monocytes (Fig. 7B). Reduced expression of *TMEM176A* and *TMEM176B* was confirmed via ddPCR of PBMC RNA (Fig. 7C, D). The levels of *TMEM176A* and *TMEM176B*, as detected by ddPCR, were tightly associated with one another, and a subset of *MAPT* pathogenic variant carriers (5 of 8) displayed lower expression of both genes than any non-carrier controls (Fig. S5D).

**Figure 7.**
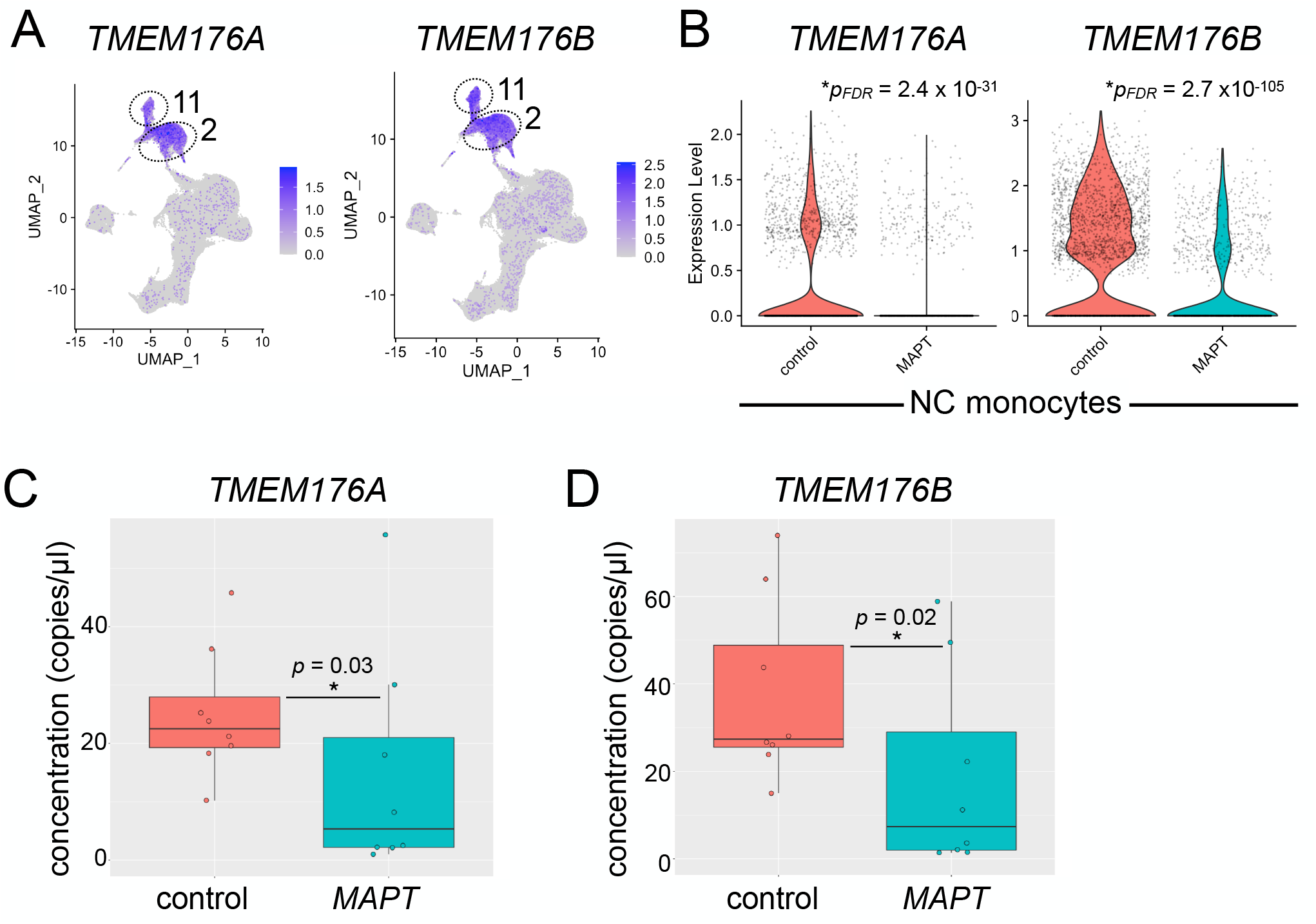
Analysis of *TMEM176A*/*B* in *MAPT* pathogenic variant carriers. **A** *TMEM176A*/*B* are highly expressed in both classical (cluster 2) and NC (cluster 11) monocytes. **B** *TMEM176A*/*B* are significantly reduced in NC monocytes (*TMEM176A, p*_*FDR*_ = 2.4 × 10^−31^; *TMEM176B, p*_*FDR*_ = 2.7 × 10^−105^) from *MAPT* carriers. These genes were similarly reduced in *MAPT* carrier classical monocytes (cluster 2; Table S3). The reduction in *TMEM176A* (**C**) and *TMEM176B* (**D**) in *MAPT* variant carriers was confirmed using bulk PBMC RNA and ddPCR (*TMEM176A, p* = 0.03; *TMEM176B, p* = 0.02).

Finally, we examined the expression of *C3AR1* given its importance in models of tauopathy [67]. *C3AR1* expression was enriched in NC monocytes (Fig. S6A) and was strikingly reduced in this cell type in *MAPT* variant carriers (Fig. S6B). *C3AR1* expression, as measured by ddPCR of PBMC RNA, trended toward reduction but did not achieve significance (Fig. S6C). However, given the importance of this gene and the complement pathway more generally in tauopathy and AD, analysis of peripheral *C3AR1* expression in larger cohorts is still warranted.

## Discussion

This study represents an effort to discover novel, blood-based biomarkers of tauopathy, and– more broadly–to begin to understand the nature of the peripheral leukocyte response to primary tauopathy. To this end, we conducted an unbiased scRNA-seq survey of ∼181,000 PBMCs in carriers of pathogenic *MAPT* variants and healthy, non-carrier controls. In doing so, we uncovered novel, peripheral blood transcriptomic signatures of familial tauopathy at single-cell resolution. In particular, we observed a significant reduction in circulating NC monocytes and numerous DEGs that were particularly enriched in specific myeloid and NK cell clusters. Next, we validated changes in several candidate DEGs selected on the basis of plausible biological relevance. These included *CX3CR1, FCGR3A*, and *TMEM176A/B*. In addition, *C3AR1*, although not found to be significantly reduced in *MAPT* carriers by ddPCR, still trended toward reduced expression.

The differential expression of *CX3CR1* observed by scRNA-seq was not only replicated via ddPCR analyses but also confirmed using a publicly available mouse microglia bulk RNA-seq dataset derived from the *MAPT* P301S model. This suggests *CX3CR1* may have potential utility as a peripheral biomarker of tauopathy and warrants further study in larger cohorts. Intriguingly, mouse *Cx3cr1* has been known for over a decade to control the levels of NC monocytes [71–73]. In addition, mouse *Cx3cr1*-mediated control of NC monocyte levels has more recently been suggested to modulate the innate immune response to traumatic brain injury [74]. Collectively, these findings suggest that the reduction in *CX3CR1* expression we observe in *MAPT* pathogenic variant carrier NC monocytes (on a per-cell basis) may be directly related to the concomitant reduction in NC monocyte abundance also observed in *MAPT* carriers.

Quite aside from the role of *Cx3cr1* in controlling circulating NC monocyte levels, *Cx3cr1* exerts a well-established, microglia-mediated modulatory effect on tau neuropathology in mouse models [57–59,75–81]. In particular, deletion of *Cx3cr1* promotes hallmark neuropathological features of tauopathy including tau hyperphosphorylation and aggregation [57–59,81]. More-recent studies on induced pluripotent stem cell-derived microglia-like cells have confirmed the importance of *CX3CR1* in regulating human microglial homeostasis [82], consistent with *Cx3cr1*’s established role in promoting the homeostatic microglial phenotype in mice [83]. Given that microglial homeostasis is dysregulated in tauopathy [84–87], our novel findings–coupled with the well-defined relationship between *CX3CR1* and tauopathy–provide a promising foundation for further investigation of this gene as a peripheral biomarker of tauopathy.

As noted above, we observed a significant reduction in NC monocytes in *MAPT* carriers relative to non-carrier controls. NC monocytes are recruited to sites of vascular damage, infection, or inflammation to patrol the local environment [73]. At these compromised sites, chemoattractant factors are released, and NC monocytes respond through the expression of cognate receptors, including CX3CR1 [73]. Strikingly, NC monocytes are thought to be reduced in peripheral blood in a variety of neurodegenerative diseases, including ALS [15,16], PD [14], and the adult-onset, hereditary leukoencephalopathy, ALSP [17]. Conversely, NC monocyte levels may be increased in the CSF in PD, suggesting a possible shift of this monocyte population from blood to CSF in the context of neurodegeneration [88]. Considering these findings, and given that: (i) ALS exists along a spectrum with FTD [89]; (ii) a portion of PD risk is mediated by variation near the *MAPT* locus [90,91]; and (iii) ALSP can manifest clinically as FTD [18], our finding of reduced NC monocytes in familial tauopathy both strengthens and extends the purported relevance of this monocyte population in neurodegenerative disease.

We observed significantly reduced expression of the canonical NC monocyte marker gene *FCGR3A* as well as significant alterations in two additional NC monocyte marker genes (*VMO1* and *IFITM3*) in *MAPT* pathogenic variant carriers by scRNA-seq. Given that *FCGR3A* expression was reduced not only in NC monocytes but also in NK cells and considering that NC monocyte abundance was simultaneously reduced, we reasoned that bulk assessment of PBMC RNA via ddPCR would be well suited to detect reduced expression of *FCGR3A* in *MAPT* carriers, and, indeed, this is what we found. This cellular phenotype suggests several possibilities. First, the reduced abundance of NC monocytes and diminished expression of *FCGR3A* on the remaining NC monocytes could reflect migration of mature NC monocytes (that express the highest levels of *FCGR3A*) out of the blood and into another compartment (e.g., CSF or brain). Second, our findings could reflect impaired survival of NC monocytes in tauopathy. Third, lower *FCGR3A* levels might reflect impaired differentiation of NC monocytes from classical or intermediate monocytes (the latter of which express intermediate levels of *FCGR3A*). We favor the first two possibilities, as the latter scenario would be expected to involve accumulation of other classes of monocytes, which we did not observe.

Our differential expression analyses identified cDCs (cluster 14) as the cell type with the highest number of DEGs with absolute LFC > 0.2. Although we did not focus on cDCs for our validation studies, we did find that normalizing NC monocyte abundance to cDC abundance enabled a clear separation of *MAPT* carriers from non-carrier controls. This finding begets the question: what is the biological link connecting NC monocytes to cDCs? Emerging literature has highlighted several intriguing connections between NC monocytes and particular subsets of DCs. For example, a putative DC subpopulation, identified transcriptomically and originally termed DC4 [92], is now considered to probably represent a subset of NC monocytes rather than DCs [46,93,94]. In addition, pathogenic variants in *STAT3* have revealed this gene’s role in regulating the production of both NC monocytes and the less-abundant cDC population, cDC1 [95]. Although we could detect the rare cDC1 population in our dataset, it became apparent only upon myeloid cell re-clustering (myeloid subcluster 7, marked by *CLEC9A* expression), and these cells were too sparse to enable us to accurately gauge their abundance or use for normalization purposes. On the other hand, it remains unclear precisely how NC monocytes are biologically related to the more-abundant cDC2 population. Nevertheless, normalizing NC monocyte abundance to either cDC2 or total cDC abundance enabled a clear separation of *MAPT* variant carriers from healthy controls. Future studies in larger cohorts will be required to determine the precise quantification metric for NC monocytes that best differentiates *MAPT* carriers from controls. It will also be important to establish whether this finding extends to sporadic forms of tauopathy; this seems likely given that similar phenomena have been reported in disparate neurodegenerative diseases.

Beyond myeloid cells, our work also highlights a potentially novel role for NK cells in primary tauopathy. In particular, NK cells had a large number of DEGs with LFC > 0.2, and our findings implicating *CX3CR1* expression not only in NC monocytes but also in NK cells as a candidate peripheral biomarker of familial tauopathy is complemented by recent research suggesting an important yet previously unappreciated role for NK cells in a mouse model of AD [96]. In addition, NK cell recruitment to the central nervous system (CNS) has been observed in ALS as well as ALS models [97]. Collectively, the available data suggest a detrimental role for NK cells in ALS and AD. More broadly, in the experimental autoimmune encephalomyelitis model of multiple sclerosis, NK cell migration into the CNS is mediated in part by CX3CR1-dependent recruitment [98,99], suggesting that the differential expression of *CX3CR1* in NK cells that we observed in our study could plausibly affect NK cell recruitment to the CNS in primary tauopathy.

Reliable biomarkers can improve diagnostic acumen and enable elucidation of specific forms of neuropathology underlying clinical dementia syndromes. For example, examination of brain structure and function via neuroimaging is a powerful method for the determination of neurodegenerative disease etiology. The use of positron emission tomography (PET) imaging, in particular, with radiotracers that bind to aggregated forms of tau has facilitated the in vivo detection of tau neuropathology in individuals with AD (reviewed in [19,100,101]). However, tau-PET tracers do not bind strongly to most forms of FTLD-tau pathology and may exhibit off-target binding in individuals with FTLD-TDP pathology [100,101]. Alternatively, the use of CSF- and blood-based protein biomarkers holds great promise for AD [19,20,102] and FTD [21,103], although in the case of FTD, we still cannot discriminate between underlying FTLD-tau and -TDP pathology. Important limitations apply to several of these methods. In particular, PET imaging is costly and available only at specialized medical centers, and CSF collection requires invasive lumbar puncture. In contrast to these methods, peripheral blood biomarkers are easy to collect and, when coupled with analytic techniques such as ddPCR, may eventually enable low-cost alternatives to today’s better-developed biomarkers.

Despite numerous advances described above, our study has several important limitations. First, due to the significant expense of scRNA-seq and our desire to capture a relatively large number of PBMCs per individual, we necessarily used a small cohort for this study. We also opted to confirm gene expression findings via an orthogonal technique using bulk PBMC RNA measurements in essentially the same cohort that was used for the scRNA-seq analysis. While we employed this strategy to minimize the possibility that technical artifacts drove discovery of the candidate genes we characterized, it will be important to evaluate the generalizability of our findings in larger cohorts. As alluded to above, it will also be important to determine which peripheral immune changes are conserved between familial tauopathy and diverse forms of sporadic primary and secondary tauopathy. In addition, given the complex temporal trajectories of brain myeloid responses in tauopathy [104], future research on large cohorts of presymptomatic *MAPT* pathogenic variant carriers will be needed to determine which peripheral changes observed here occur prior to disease onset. Finally, it will be important to ascertain whether the peripheral leukocyte changes discovered here are reflected by parallel changes in brain myeloid cells in individuals with tauopathy.

## Conclusions

To our knowledge, this is the first scRNA-seq study of peripheral blood cells in primary tauopathy. Beyond our initial discoveries, we validated a handful of DEGs via an orthogonal technique, ddPCR. In particular, we have connected longstanding observations from mouse models regarding microglial *Cx3cr1* and tau neuropathology to reduced *CX3CR1* in peripheral leukocytes in individuals with familial tauopathy. Moreover, we discovered a significant reduction in the abundance of circulating NC monocytes, a cell type that is similarly reduced in several additional neurodegenerative diseases. We also discovered large numbers of DEGs in NK cells, including *CX3CR1*, which is thought to be involved in recruitment of NK cells to the CNS. Further studies are now required to investigate the generalizability of our findings through replication in larger cohorts and extension to other tauopathies and related neurodegenerative diseases. Analogous studies of PBMCs in *GRN* pathogenic variant carriers and *C9orf72* hexanucleotide repeat expansion carriers should enable the discovery of peripheral biomarkers of FTLD-TDP. Ultimately, comparative studies should clarify the role of peripheral immune responses in distinct proteinopathies and enable discovery of novel peripheral biomarkers that can successfully discriminate between tau and TDP-43 neuropathology, providing critical new tools for diagnostics and clinical trials.

## List of abbreviations

AD: Alzheimer’s disease
ALS: amyotrophic lateral sclerosis
ALSP: adult-onset leukoencephalopathy with axonal spheroids and pigmented glia
CAT: Center for Advanced Technology
cDC: conventional dendritic cell
CDR-SB: Clinical Dementia Rating scale Sum of Boxes
CNS: central nervous system
CSF: cerebrospinal fluid
dd: droplet digital
DEG: differentially expressed gene
FDR: false discovery rate
FTD: frontotemporal dementia
FTLD: frontotemporal lobar degeneration
IHG: Institute for Human Genetics
LFC: log_2_ fold-change
LPS: lipopolysaccharide
MAC: Memory and Aging Center
MCL: Markov cluster algorithm
NC: nonclassical
NK: natural killer
PBMCs: peripheral blood mononuclear cells
PCA: principal component analysis
PD: Parkinson’s disease
PET: positron emission tomography
QC: quality control
RIN: RNA integrity number
RT: reverse transcription
scRNA-seq: single-cell RNA sequencing
UCSF: University of California, San Francisco
UMAP: uniform manifold approximation and projection

## Declarations

### Ethics approval and consent to participate

All participants or their surrogates provided written informed consent prior to study participation; all aspects of the studies described here were approved by the UCSF institutional review board.

### Consent for publication

Not applicable.

### Availability of data and materials

The scRNA-seq dataset described here will be uploaded to the FAIR Data Sharing Portal within the Alzheimer’s Disease Workbench, which is supported by the Alzheimer’s Disease Data Initiative, and can be accessed at https://www.alzheimersdata.org/ad-workbench.

### Competing interests

JSY serves on the scientific advisory board for the Epstein Family Alzheimer’s Research Collaboration. ALB has served as a consultant for Aeovian, AGTC, Alector, Arkuda, Arvinas, AviadoBio, Boehringer Ingelheim, Denali, GSK, Life Edit, Humana, Oligomerix, Oscotec, Roche, Transposon, TrueBinding and Wave.

### Funding

Funding to JSY and for this project comes from NIH-NIA K01AG049152, R01AG062588, R01AG057234, P30AG062422, P01AG019724; NIH-NINDS U54NS123985; the Rainwater Charitable Foundation; the Larry L. Hillblom Foundation; the Bluefield Project to Cure Frontotemporal Dementia; the Alzheimer’s Association; the Global Brain Health Institute; the French Foundation; and the Mary Oakley Foundation. VES is supported by R01AG052496. SEL is supported by R01AG058233. ALB receives research support from NIH (U19AG063911, R01AG038791, R01AG073482), the Tau Research Consortium, the Association for Frontotemporal Degeneration, Bluefield Project to Cure Frontotemporal Dementia, Corticobasal Degeneration Solutions, the Alzheimer’s Drug Discovery Foundation, and the Alzheimer’s Association, and has received research support from Biogen, Eisai and Regeneron. The content of this publication is solely the responsibility of the authors and does not necessarily represent the official views of the National Institutes of Health.

### Authors’ contributions

DWS initiated the study, analyzed the scRNA-seq and ddPCR data, and wrote the manuscript. CWS analyzed the scRNA-seq data and contributed to writing the manuscript. TPJ isolated PBMCs and performed ddPCR experiments with DWS. LWB analyzed scRNA-seq data and edited the manuscript. VES, SEL, KPR, HJR, ALB, WWS, and BLM were involved in research participant recruitment and clinical characterization. EGG was involved in early stages of the scRNA-seq analysis and provided guidance on study design. JSY initiated the study, oversaw the analysis, and edited the manuscript.

## Acknowledgements

We thank the UCSF Genomics CoLab and IHG staff, including Catherine Chu, for expert technical assistance. In addition, we thank UCSF CAT staff, including Tyler Miyasaki, for expert advice on the use of ddPCR equipment. The cartoon in Figure 4D was created with BioRender.com.

## Figure Legends

**Figure S1.**
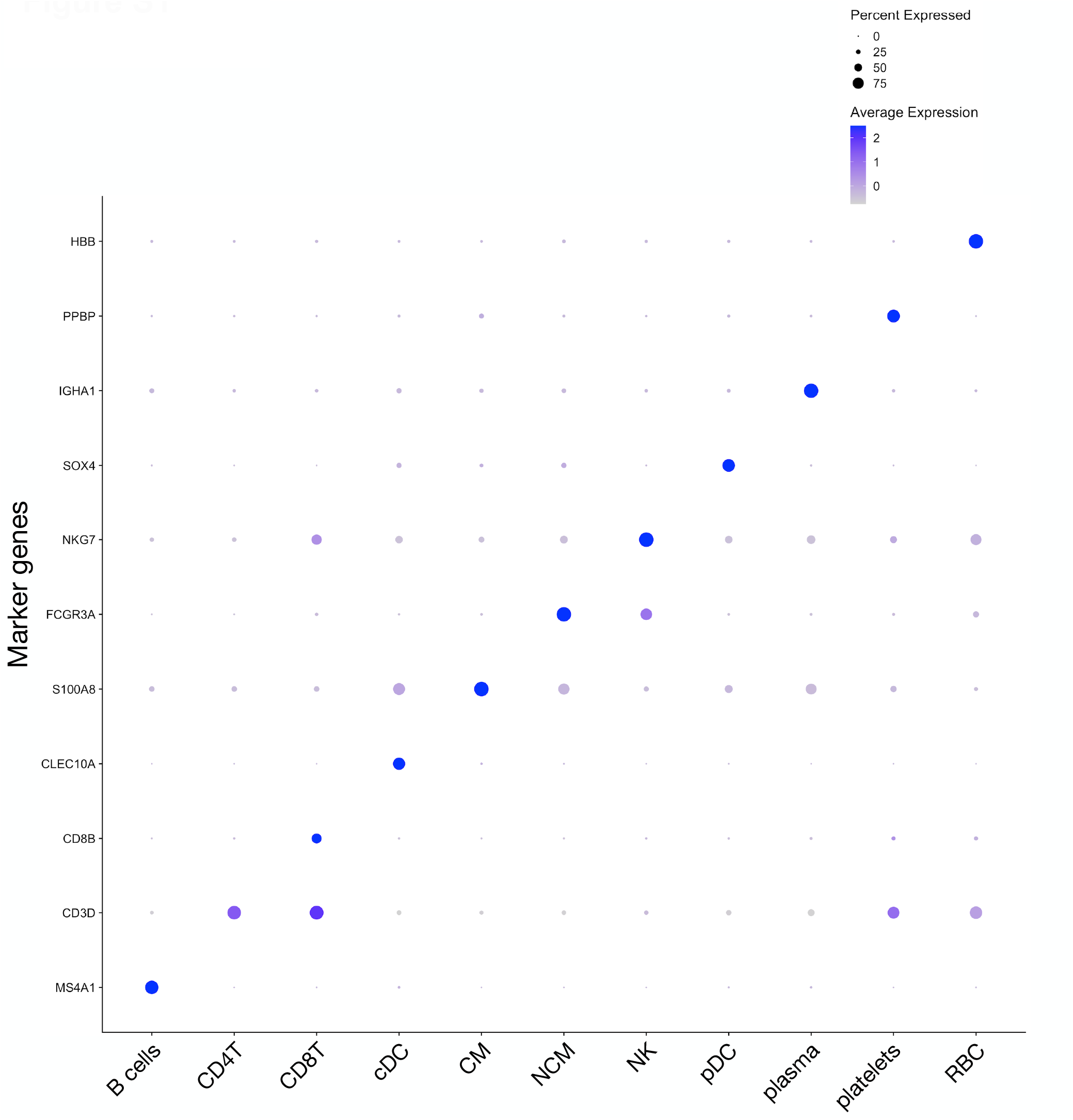
Annotation of cell types from cluster markers. Cell types were annotated using a combination of canonical marker genes and additional markers identified via Seurat’s `FindMarkers` function. Clusters were assigned to the following cell-type categories: B cells (clusters 8, 12, 19), CD4^+^ T cells (CD4T; clusters 0, 1, 7, 10, 17), CD8^+^ T cells (CD8T; clusters 4, 5, 6, 9), cDC (cluster 14), classical monocytes (CM; cluster 2), NC monocytes (NCM; cluster 11), NK cells (clusters 3 and 15), plasmacytoid dendritic cells (pDC; cluster 16), plasma cells (cluster 18), and contaminating platelets (cluster 13) and red blood cells (RBC; cluster 20). Expression of marker genes is shown using Seurat’s `DotPlot` visualization, with dot size indicating the percentage of cells in a given category having detectable expression of a given marker gene and color indicating the average expression level.

**Figure S2.**
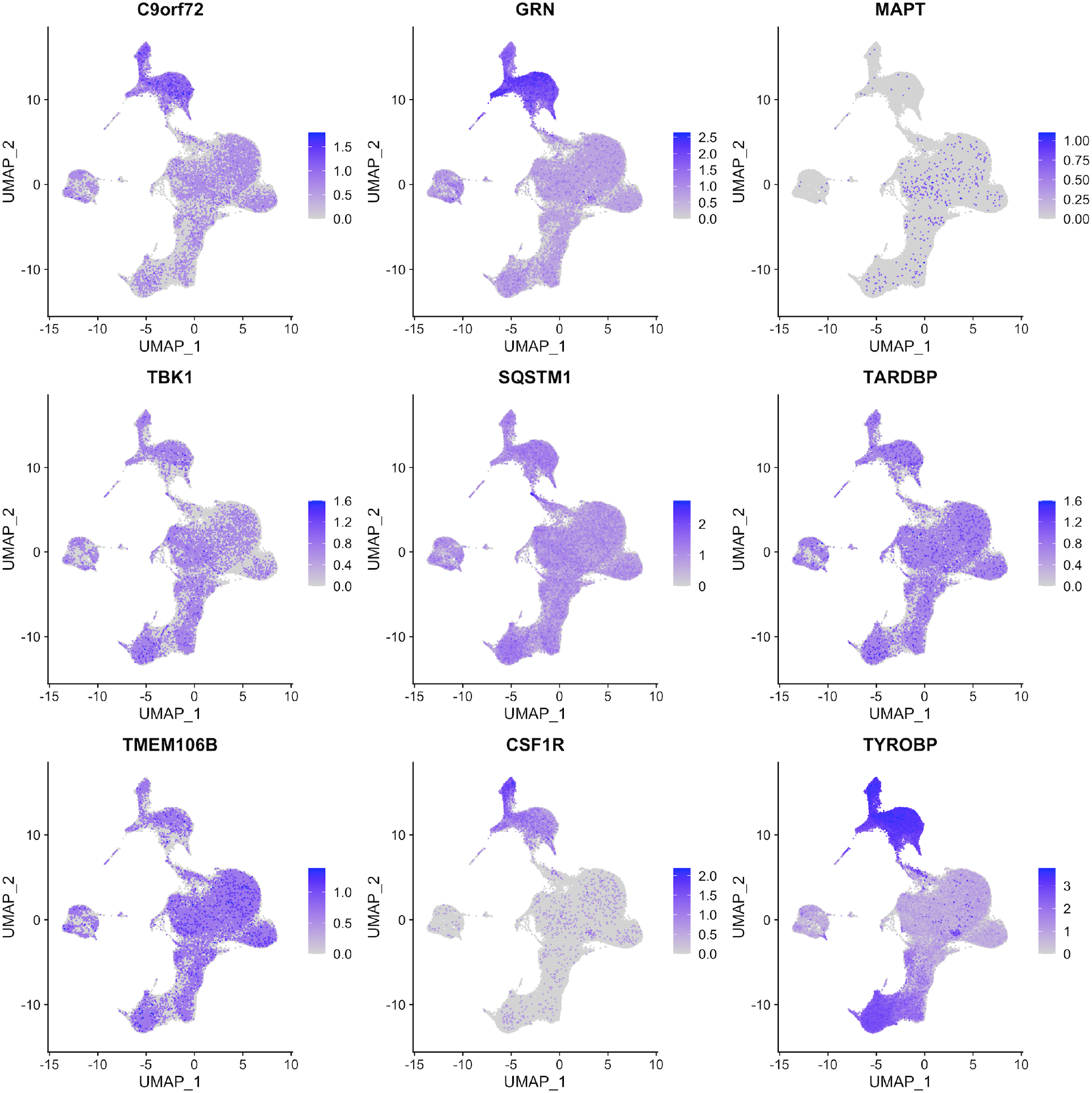
Expression of FTD-associated genes in PBMCs. The expression level of nine FTD-associated genes (*C9orf72, GRN, MAPT, TBK1, SQSTM1, TARDBP, TMEM106B, CSF1R, TYROBP*) across all PBMCs in the dataset is depicted. Some genes displayed widespread expression across many PBMC cell types (*TBK1, SQSTM1, TARDBP, TMEM106B*), while others showed enriched expression in myeloid clusters (*C9orf72, GRN, CSF1R, TYROBP*). In contrast to other FTD-associated genes, *MAPT* expression was very rarely detected in PBMCs.

**Figure S3.**
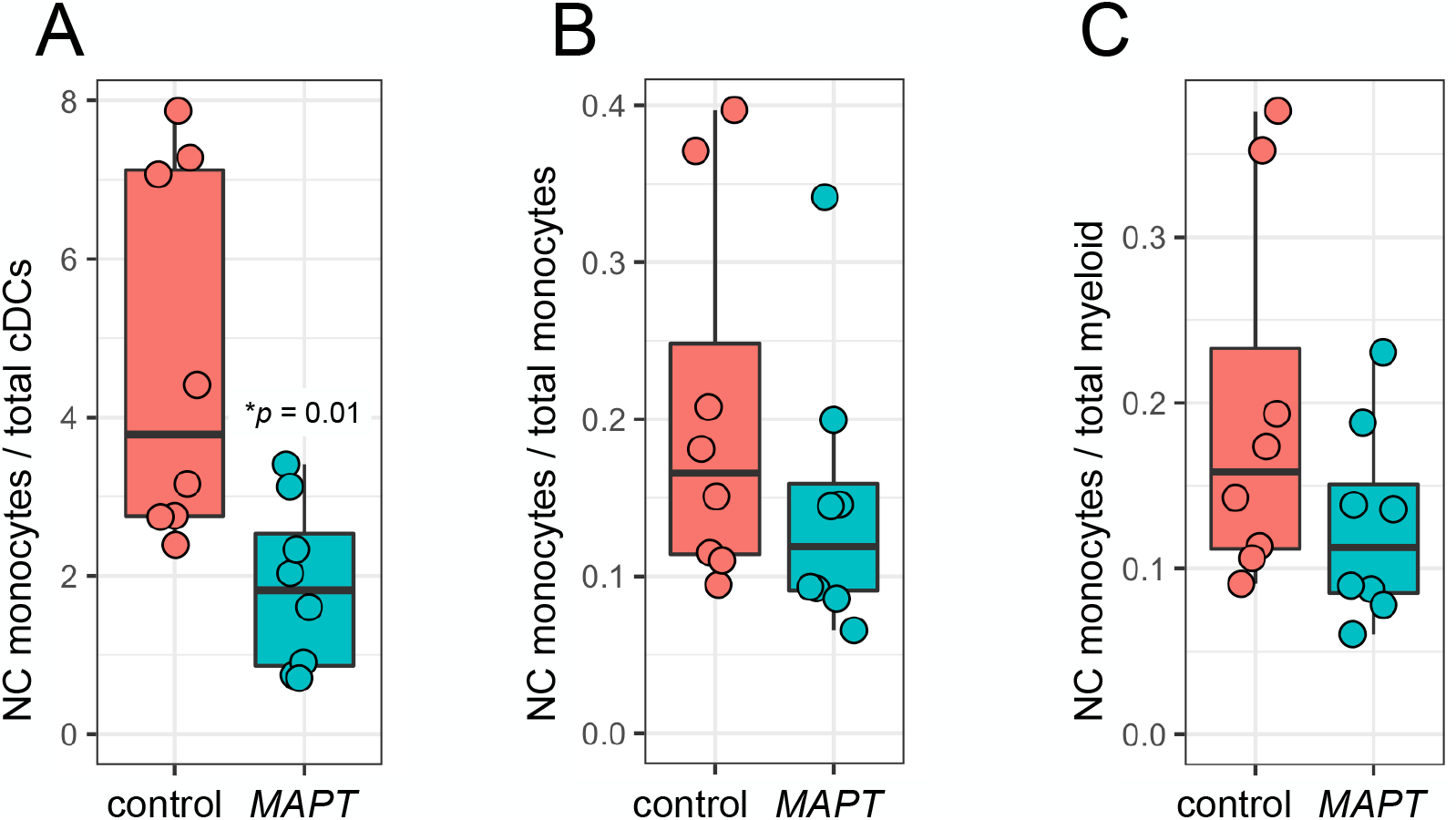
Nonclassical monocyte abundance relative to other myeloid populations. **A** *MAPT* pathogenic variant carriers showed a significant reduction (*p* = 0.01) in the ratio of NC monocytes to total cDCs (cDC1 + cDC2). **B, C** In contrast, the ratio of NC monocytes to total monocytes (**B**) or total myeloid cells (**C**) was not significantly reduced in *MAPT* variant carriers compared to non-carrier controls.

**Figure S4.**
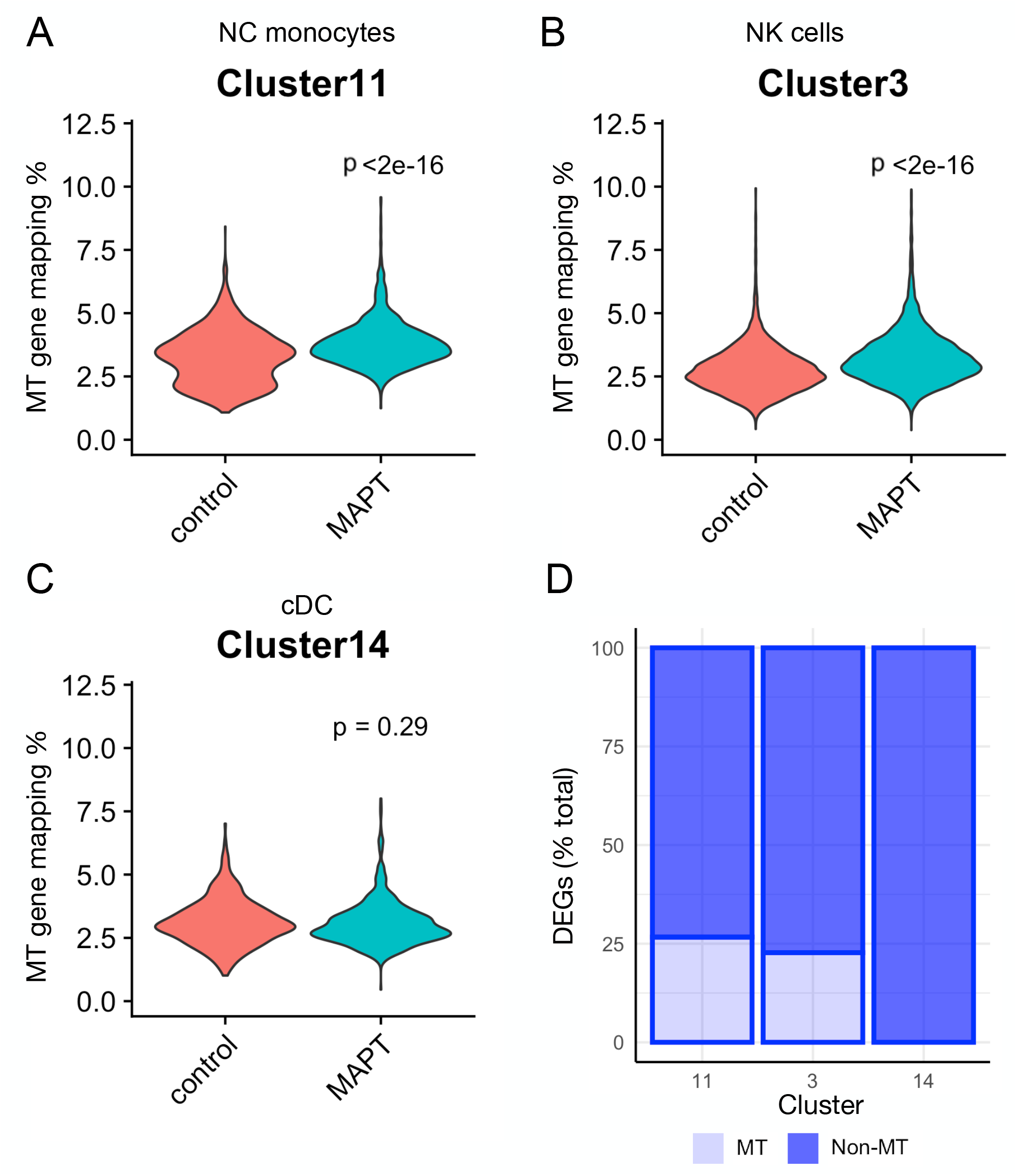
Mitochondrial genome mapping percentages and mitochondrial DEGs in selected clusters. **A, B** Mitochondrial mapping percentage was subtly but significantly increased (*p* < 2 × 10^−16^) in *MAPT* pathogenic variant carriers in clusters 11 (NC monocytes; **A**) and 3 (NK cells; **B**). **C** On the other hand, mitochondrial mapping percentage was not significantly increased in cluster 14 (cDC). **D** Clusters 11 and 3 both harbored appreciable fractions of mitochondrial DEGs relative to total DEGs, while cluster 14 did not contain mitochondrial DEGs.

**Figure S5.**
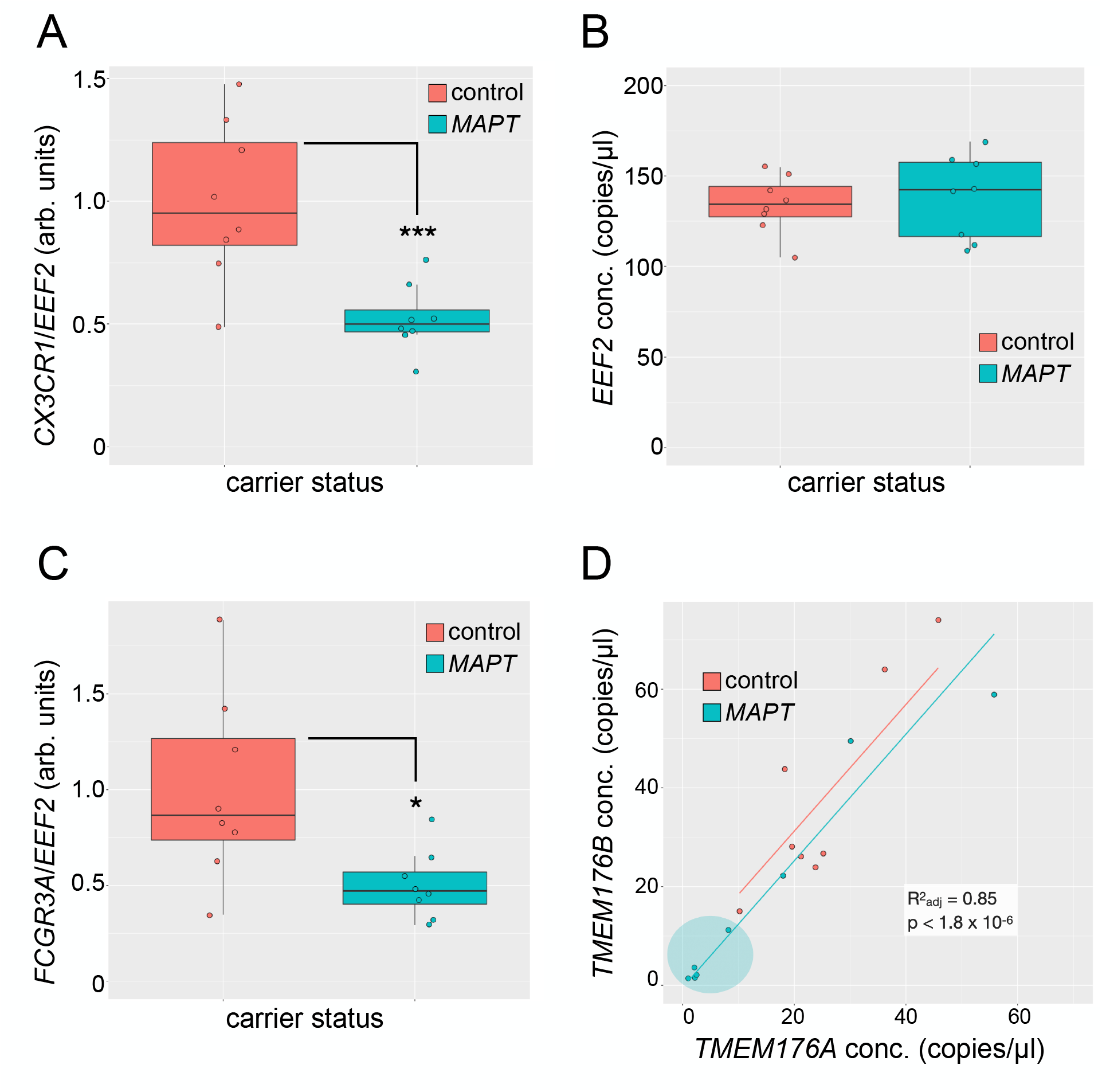
Additional droplet digital PCR analysis. **A** Normalization of the *CX3CR1* concentration to that of reference gene *EEF2* produced results similar (***, *p* = 0.0006) to what is depicted in Fig. 5A. **B** *EEF2* absolute concentration values were similar between *MAPT* pathogenic variant carriers and non-carrier controls. **C** Normalization of the *FCGR3A* concentration to that of *EEF2* did not alter the results depicted in Fig. 6B (*, *p* = 0.01). **D** *TMEM176A*/*B* levels were closely associated with one another (R^2^_adj_ = 0.85, *p* < 1.8 × 10^−6^). A subset of *MAPT* variant carriers (5 of 8) showed levels of *TMEM176A*/*B* that were lower than what was observed for any non-carrier control.

**Figure S6.**
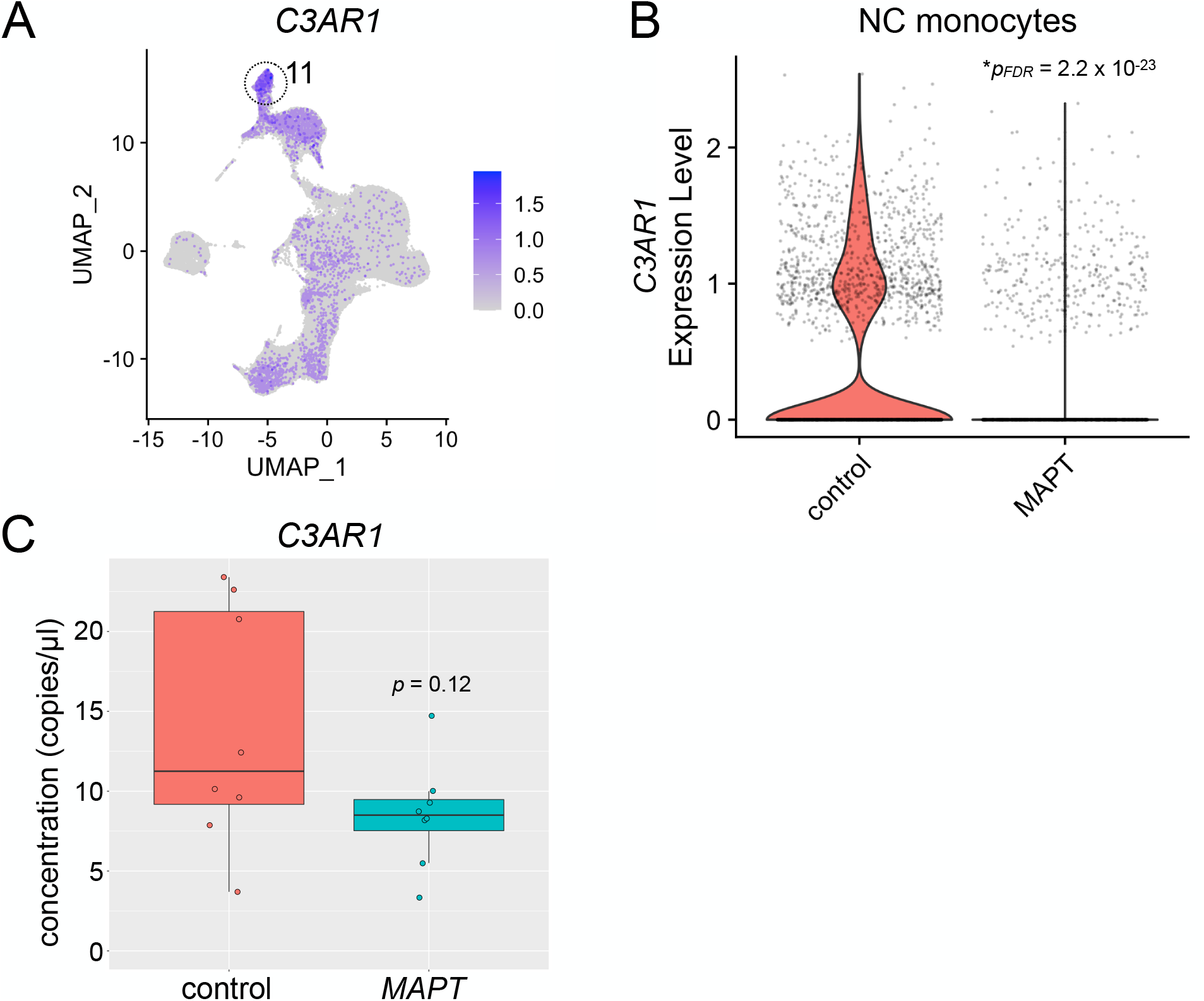
Potential dysregulation of *C3AR1* in *MAPT* pathogenic variant carriers. **A** *C3AR1* expression was enriched in the NC monocyte cluster (11). **B** *C3AR1* expression in NC monocytes was significantly reduced in *MAPT* variant carriers (\**p*_*FDR*_ = 2.2 × 10^−23^). **C** ddPCR analysis of PBMC RNA revealed a trend toward reduced expression of *C3AR1* in *MAPT* variant carriers which did not reach significance (*p* = 0.12).

